# Activated Notch1 Redirects CD4-Fated CD4^+^ CD8^+^ Precursors to the CD8 Lineage During Thymocyte Selection Without Causing T Cell Leukemia

**DOI:** 10.1101/2025.02.28.640858

**Authors:** Mahmoud El-Maklizi, Philaretos C. Kousis, Julie S. Yuan, Blair MacDonald, Mark Gower, Miki S. Gams, Amine Boudil, Cynthia J. Guidos

## Abstract

Positive selection of CD4/CD8 double-positive (DP) thymocytes generates helper and cytotoxic T lineages whose CD4 or CD8 co-receptor expression matches their αβ T cell receptor (TCRαβ) specificity MHCII or MHCI, respectively. Notch1 signaling is critical for early T cell development and was suggested to regulate CD4/CD8 lineage determination during DP thymocyte selection. However, early studies expressing activated Notch1 in pre-DP progenitors did not resolve this question. Here we showed that CD4-Cre-mediated activation of canonical *Rbpj*-dependent Notch1 signalling potently skews DP selection to the CD8 lineage. We used a tamoxifen-inducible “time-stamp” model coupled with phenotypic staging to show that that activated Notch1 decreases generation of Gata3^+^ CD4^+^CD8^lo^ selection intermediates during the earliest stages of DP selection, thereby preventing ThPOK induction and emergence of CD4 lineage cells. Finally, activated Notch1 efficiently re-directed DP thymocytes expressing MHCII-specific TCRαβ into the CD8 lineage. These studies show that activated Notch1 acts early during DP selection to skew selection of CD4-fated, MHCII-specific DP precursors to the CD8 lineage.

## INTRODUCTION

Notch1 is a transmembrane signalling receptor that plays several key roles during the early CD4/CD8 double-negative (DN) stages of intrathymic T cell development^1^. After being activated by Delta-like 4 (DL4), a Notch ligand expressed by cortical thymic epithelial cells, the intracellular region of Notch1 (ICN1) is released from its membrane tether and migrates to the nucleus where it interacts with the Rbpj transcription factor to induce target gene expression^2^. Activated Notch1 suppresses the alternative cell fates of thymus-seeding multipotent DN1 progenitors and induces T-lineage specification and commitment by the DN2 stage^3^. It then promotes clonal expansion of TCRβ^+^ DN3 progenitors to yield a large pool of DP thymocytes via a process known as beta-selection^4–7^. Although Notch1 mRNA levels decline considerably after beta-selection^8^, Notch1 protein is readily detectable on DP thymocytes by both antibody staining^9^ and by binding of soluble DL4^10^. Thus, Notch1 signalling also has the potential to influence TCRαβ-induced positive selection^11,12^ and/or differentiation of DP thymocytes into mature CD4 helper or CD8 cytotoxic single-positive (SP) lineages. However, the impact of activated Notch1 on DP thymocytes selection and CD4/CD8 lineage determination remain poorly defined.

Early studies addressed this question by expressing ICN1, which constitutively activates Notch1 in a ligand-independent fashion, in hematopoietic stem cells (HSCs) or DN2-DN3 thymocytes and then assessing their generation of mature CD4 and CD8 single-positive (SP) thymocytes. Robey et al. found that transgenic expression of ICN1 under control of the *Lck* proximal promoter (*LckPr*), which is highly active beginning at the DN2 stage, promoted the positive selection of CD8 SP at the expense of CD4 SP thymocytes^13^. Interestingly, *LckPr-ICN1* promoted the selection of mature CD8 SP thymocytes in mice lacking MHC class I (MHCI), but not in mice that also lacked MHC class II (MHCII). These data suggested that activated Notch1 could re-direct DP progenitors expressing MHC II-specific TCRαβ, which are normally CD4-fated, into the CD8 SP lineage. Surprisingly, a different *LckPr-ICN1* transgene promoted survival and development of both CD4 and CD8 SP lineages independently of MHC recognition^14^. In a third study, DP thymocytes derived from *ICN1-*transduced HSC showed impaired TCRαβ signalling and positive selection^15^. Collectively, these early studies showed that Notch1 activation profoundly affects positive selection and CD4/CD8 lineage determination of DP thymocytes, albeit in model-specific ways.

Ectopic activation of Notch1 in HSC or DN2/DN3 thymocytes also induces the development of T cell lymphoblastic lymphomas with high penetrance^15–19^, presenting challenges with using these models to study Notch1 functions in later stages of T cell development. To bypass this problem, other groups used *in vitro* approaches to manipulate Notch1 activation specifically in DP thymocytes. One study found that genetic or antibody-mediated inhibition of Notch1 signalling blocked *in vitro* differentiation of CD8 but not CD4 SPs^20^. However, CD4Cre-induced inactivation of Notch/Rbpj signaling in polyclonal DP thymocytes does not impact generation of conventional CD4 or CD8 SP cells^21,22^. Nonetheless, ligand-induced Notch activation is necessary for *in vitro* selection of DP thymocytes expressing certain MHCI-restricted TCRαβ into the CD8 lineage^23^. Thus, physiological Notch/Rbpj signaling may only impact selection of cells with particular TCR signaling strengths.

Recent studies have identified the key cellular and molecular sequences that govern DP selection and CD4/CD8 lineage commitment. Selection begins when DP thymocytes are signaled by self-peptide MHC II (pMHCII) or pMHC-I complexes expressed by thymic epithelial cells in the thymic cortex to upregulate CD69 and Gata3^24,25^. This transcription factor is required to generate CD4^+^CD8^lo^ selection intermediates^26^, which undergo CD4/CD8 lineage determination in response to differential signalling by TCR-pMHCI vs TCR-pMHCII^27–30^. CD8 downregulation does not disrupt TCR-pMHCII signalling in CD4^+^ CD8^lo^ thymocytes that express pMHCII-specific TCRαβ, sustaining Gata3 expression which induces then ThPOK^26^. The latter factor mediates CD4 lineage commitment by blocking expression of Runx3^31–33^, the CD8 lineage-determining factor^34–38^. By contrast, CD8 downregulation disrupts TCR-pMHCI signalling in CD4^+^CD8^lo^ cells that express pMHCI-specific TCRαβ. Thus, these cells fail to maintain Gata3 and do not induce ThPOK, allowing upregulation of Runx3 which drives CD8 lineage commitment^39^. These early selection intermediates express CCR9 but not CCR7, retaining them in the thymic cortex by CCR9 interaction with its ligands^40,41^. However, as signalled DP thymocytes mature they upregulate CCR7^42,43^ and eventually lose CCR9 and migrate to the thymic medulla to negative selection.

It remains unclear how activated Notch impacts CD4/CD8 lineage determination and the distinct stages of DP selection. Furthermore, the interplay between Notch1 signalling and the molecular determinants of the CD4/CD8 lineage decision has not been explored. To answer these questions, we created a CD4-Cre-induced Notch1 gain-of-function mouse model to induce ligand-independent Notch1 activation at the DP stage of thymocyte development, a strategy that prevented the development of thymic lymphoma. We also created a tamoxifen-inducible *CD4-CreER^T^*^2^ model to follow the *in vivo* fate of “time-stamped” DP thymocytes after induction of *ICN1-GFP* or *YFP*. Using these models, we showed that activated Notch1 profoundly alters the earliest stages of DP selection to promote the CD8 fate at the expense of the CD4 fate by blocking TCR-induced upregulation of Gata3 and generation of CD4^+^ CD8^lo^ selection intermediates, thus impairing subsequent ThPOK expression. Our study has resolved the long-standing uncertainty from earlier studies on how activated Notch1 affects positive selection and provides novel insights into the mechanisms involved.

## RESULTS

### CD4-Cre-induced Notch1 activation favors selection of CD8 SP thymocytes without causing T cell leukemia

To assess the impact of Notch1 signalling on DP thymocyte selection, we created a new mouse model to express activated Notch1 beginning at the DP stage of thymocyte development. We obtained mice that contain sequences encoding *ICN1*, an internal ribosomal entry sequence (*IRES*) and *GFP* inserted into the *Rosa26 (R26)* locus^44^ and crossed them to a *CD4-Cre* transgenic strain in order to promote constitutive (Con) expression of both *ICN1* and *GFP* from the *R26* locus in CD4-expressing thymocytes. We refer to this *CD4-Cre*; *R26^ICN1-GFP^* strain as *ICN1-GFP^Con^*. We used flow cytometry to track ICN1-GFP expression and its impact on thymocyte differentiation in young adult mice. Interestingly, the percentage of all CD4/CD8 thymocyte subsets was significantly different in *ICN1-GFP^Con^*thymocytes than in their *CD4-Cre; R26^WT^* littermates (referred to as *CD4-Cre*) littermates (Fig. 1 A, B). GFP expression increased with TCRβ expression in *ICN1-GFP^Con^* thymocytes, and 70-80% of DP thymocytes were GFP^lo^ (Fig. 1 A, C). However, only a small fraction of TCRβ^-^ CD8^+^ immature SP (ISP) thymocytes expressed GFP, revealing that CD4-Cre was not highly active in these immediate precursors of DP cells^45^.

**Figure 1:**
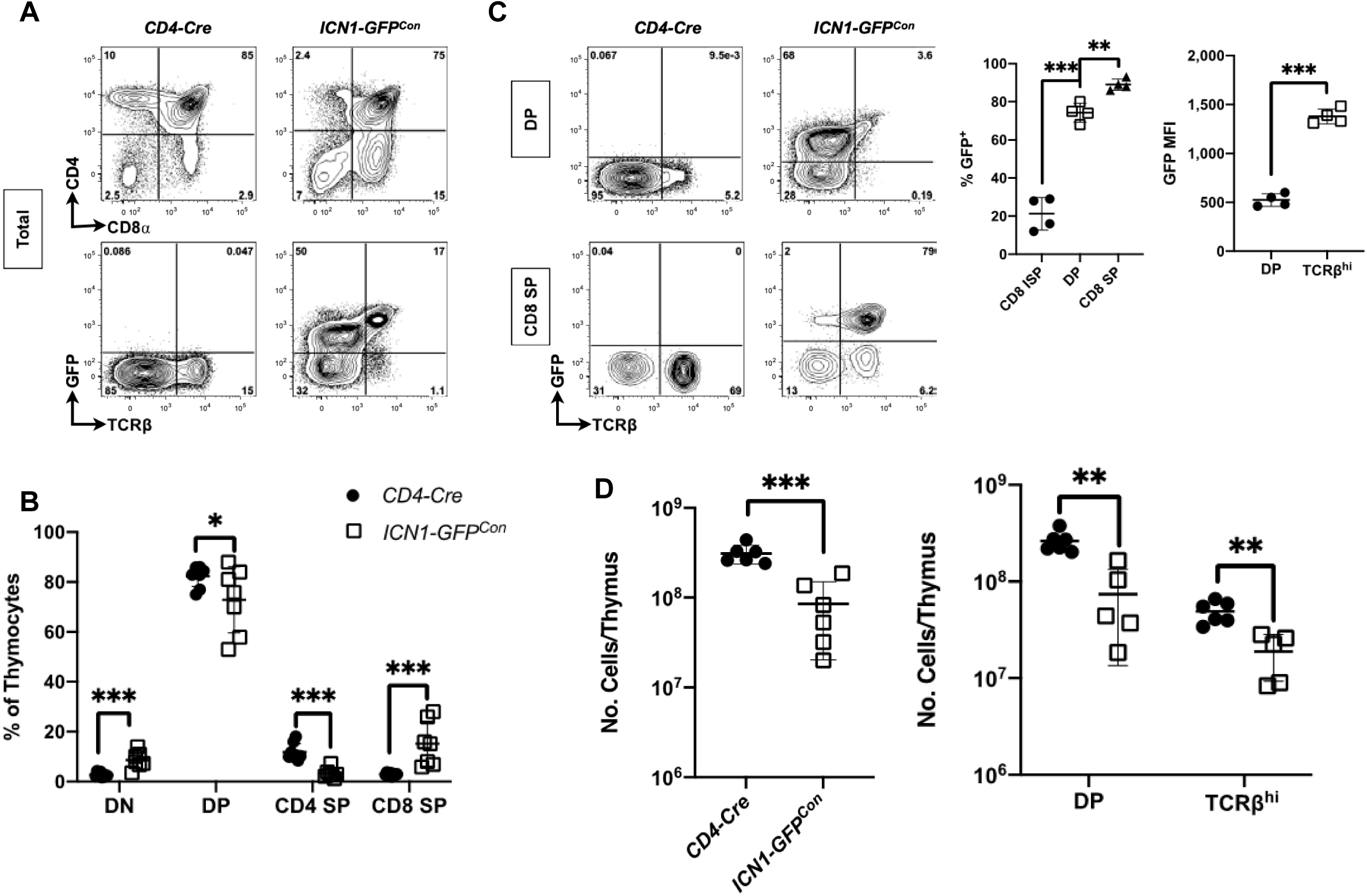
Characterization of *ICN1-GFP^Con^* thymocytes. Flow cytometric analysis of young adult thymocytes from *ICN1-GFP^Con^*and *CD4-Cre* littermate control mice. Data were pooled from 3 experiments with n=4-8 per genotype. (A) Representative flow cytometry contour plots of CD4 versus CD8α and TCRβ versus GFP gated on total live single cells from each strain. The percentage of cells in each quadrant is shown. (B) Scatter dot plot shows the percentage of each subset by strain: DN: CD4^-^ CD8α^-^, DP: CD4^+^ CD8α^+^, CD4 SP: TCRβ^+^ CD4^+^ CD8α^-^ and CD8 SP: TCRβ^+^ CD4^-^ CD8α^+^ thymocytes. Multiple *t* tests: *, *q*=0.02; ***, *q*=<0.001. (C) ICN1-GFP expression increases during thymocyte differentiation. Representative contour plots of TCRβ versus GFP gated on DP or total CD8 SP thymocytes from each strain. Scatter dot plots show: *Middle:* the percentage of GFP^+^ cells among CD8 ISP (TCRβ^-^ CD4^-^ CD8α^+^), DP and CD8 SP (TCRβ^+^ CD4^-^ CD8α^+^) thymocytes. Post-hoc *t* tests compared each subset compared to DP cells: **, *q*=0.007; ***, *q*<0.001. *Right:* GFP MFI on DP and TCRβ^hi^ thymocytes in *ICN1-GFP^Con^* mice. Unpaired 2-tailed *t* test: ***, *p*<0.001. (D) Reduced abundance of DP and TCRβ^hi^ thymocytes in *ICN1-GFP^Con^* mice. Dot plot shows the total thymic cellularity (*Left*, ***, *p*<0.001) and the number of DP and TCRβ^hi^ thymocytes (*Right,* multiple t-tests, **, *q*=0.002) in each thymus.

We performed global gene expression profiling of sorted DP thymocytes each strain and showed that canonical Notch target genes, such as *Dtx1, Hes1, Fjx1, Ptcra* and including *Notch1* itself, were expressed at significantly higher levels in DP thymocytes from *ICN1-GFP^Con^* mice, confirming that Notch1 signalling was strongly activated (Suppl. Table 1A). However, other genes identified as Notch1 targets in more immature T cells (*Nrarp*^46–48^*, Il2ra*^4,14^*, Tcf7*^49^ and *Lef1*^50^) or T cell leukaemias (*Il7r, Depdc6*) were not differentially expressed (Suppl. Table 1B), suggesting context-dependent activation of Notch1 targets during T cell development. In addition, neither *Myc* nor several other targets implicated in Notch1-induced T cell leukemogenesis^51,52^ were not upregulated by constitutive Notch1 activation in DP thymocytes (Suppl. Table 1C). Indeed, mutants exhibited 32% lower thymic cellularity, reflecting a significantly lower abundance of both DP and mature TCRβ^hi^ thymocytes, than *CD4-Cre* mice (Fig. 1D). This finding is likely explained by down-regulation of cell cycle-associated genes and pathways in mutant DP thymocytes (Suppl. Tables 2, 3). Collectively, these data show that Notch1 was strongly activated in DP thymocytes from *ICN1-GFP^Con^* mice, but this did not promote leukemic transformation.

To examine the impact of ligand-independent Notch1 activation on thymic selection, we quantified the frequency of mature subsets among post-selection TCRβ^hi^ cells thymocytes. *ICN1-GFP^Con^* mice had significantly lower TCRβ^hi^ CD4 SP and significantly higher percentages of DN and CD8 SP thymocytes among TCRβ^hi^ cells compared to *CD4-Cre* mice (Fig. 2A). To account for the reduced precursor pool of DP thymocytes (Fig. 1D), we calculated the ratio of the number of each TCRβ^hi^ subset relative to the number of DP thymocytes in individual *ICN1-GFP^Con^*vs *CD4-Cre* mice. There was no difference in the normalized output of selected TCRβ^hi^ DP thymocytes, suggesting that *ICN1* does not impact the early selection of DP cells (Fig. 2B). In contrast, *ICN1* significantly reduced the output ratio of TCRβ^hi^ CD4 SP thymocytes and increased the output ratio of TCRβ^hi^ DN and TCRβ^hi^ CD8 SP thymocytes. Transgenic expression of *hBCL2* significantly improved DP thymocyte viability in *ICN1-GFP^Con^* mice but did not restore their generation of TCRβ^hi^ CD4 SP thymocytes (Fig. 2C). In contrast, *Rpbj* deletion in *ICN1-GFP^Con^*mice restored CD4 SP and CD8 SP frequencies to those seen in *CD4-Cre* mice (Fig. 2D). These data show that Notch1 activation alters the selection of DP thymocytes in an *Rbpj*-dependent fashion to favour the selection of mature DN and CD8 SP at the expense of CD4 SP thymocytes.

**Figure 2:**
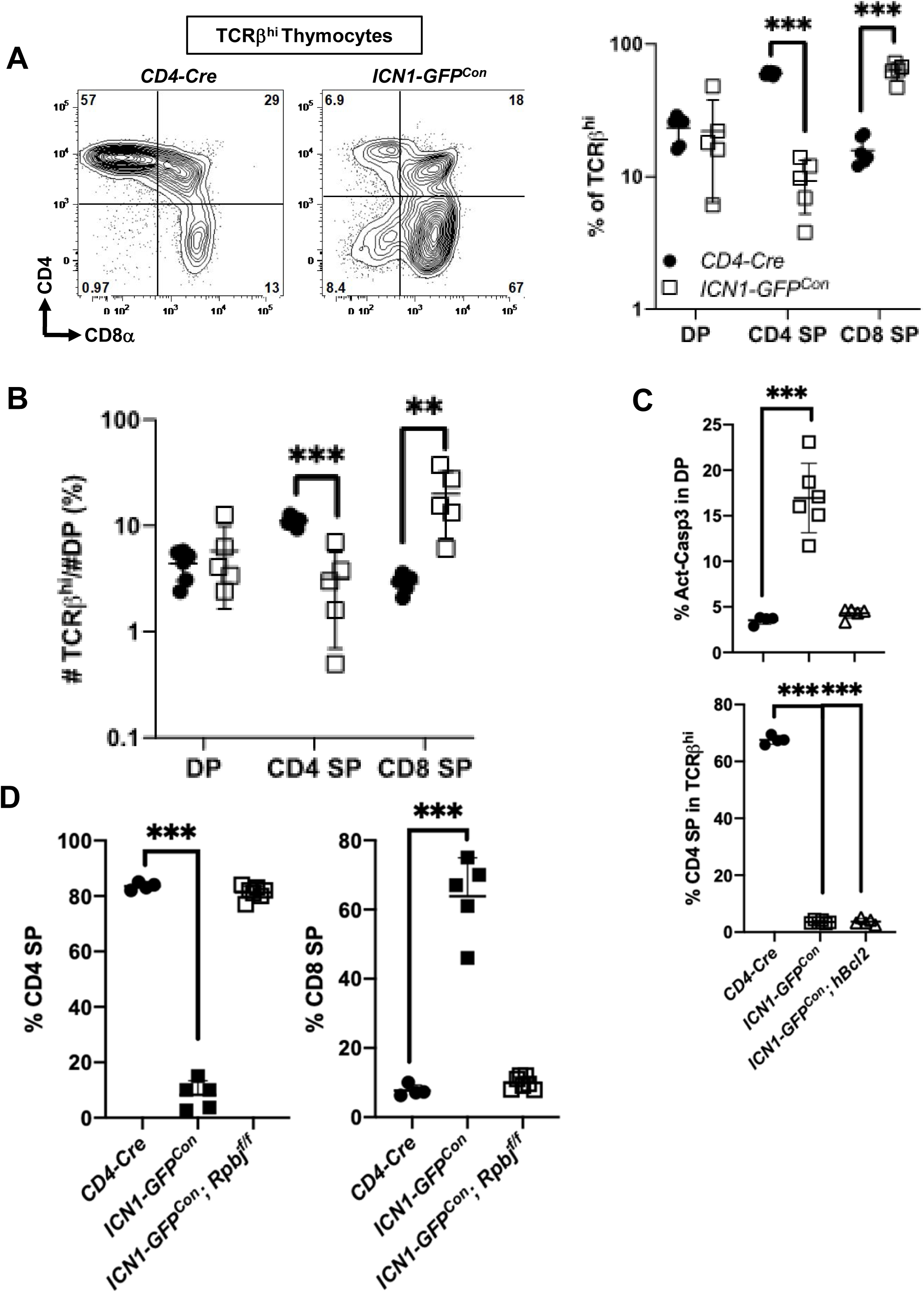
Notch1 activation in DP thymocytes skews thymic output towards the CD8 lineage. Analysis of TCRβ^hi^ thymocytes from the experiments shown in Fig. 1. (A) Representative gating of flow cytometry contour plots of CD4 versus CD8α on live single TCRβ^hi^ thymocytes in each strain. Scatter dot plot shows the percentage of DN, DP, CD4 and CD8 SP cells among TCRβ^hi^ thymocytes in each strain. Multiple *t* tests: **, *q*=0.006; ***, *q*<0.001. (B) Output ratios for each TCRβ^hi^ subset (DN, DP, CD4 SP and CD8 SP) were calculated as: (#TCRβ^hi^ subset/#DP thymocytes) x 100. Multiple *t* tests, *: *q*= 0.01; **: *q*=0.004, ***, *q*<0.001. (C) Transgenic *hBCL2* improved DP survival but did not restore CD4 SP differentiation to *ICN1-GFP^Con^* mice. Scatter dot plots show the percentage of cells staining positive for activated caspase 3 among DP thymocytes (*top*) and the percentage of CD4 SP among TCRβ^hi^ thymocytes in each strain (*bottom*). (D) CD8 lineage skewing in *ICN1-GFP^Con^* mice is *Rbpj*-dependent. Scatter dot plots show the percentage of CD4 SP and CD8 SP among TCRβ^hi^ CCR7^+^ GFP^-^ or GFP^+^ thymocytes from each strain (*left*) or CD8 SP among TCRβ^hi^ CCR7^+^ GFP^-^ or GFP^+^ thymocytes from each strain (*right*). For (C) and (D), one-way ANOVA with post-hoc *t* tests comparing each strain to *CD4-Cre*, ***: *q*<0.001.

### Tamoxifen-inducible Notch1 activation skews DP selection toward the CD8 SP lineage

We next created *R26^ICN1-GFP^* mice expressing the *CD4-CreER^T2^* transgene^53^ to allow tamoxifen-inducible (Ind) Notch1 activation in DP thymocytes. To follow the fate of wild-type DP thymocytes in the same experimental context, we also generated *CD4-CreER^T2^; R26^YFP^* (referred to as *YFP^Ind^*) mice (Fig. 3A). Since activation of *CD4-CreER^T2^* by tamoxifen is not highly efficient, this model allows induction of *ICN1-GFP* or *YFP* in a small subset of DP thymocytes and their progeny within the context of a normal thymus.

**Figure 3:**
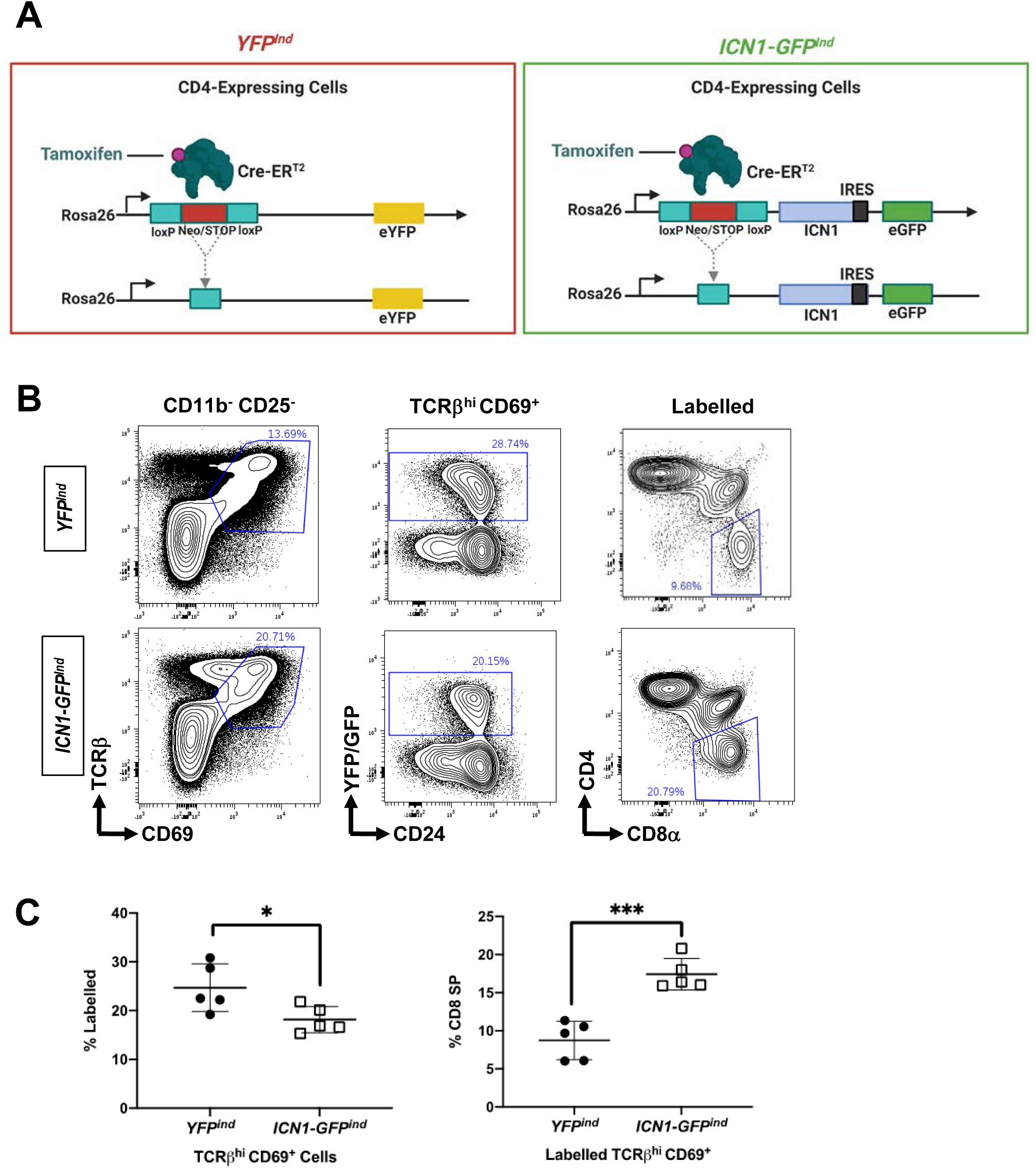
Tamoxifen-inducible Notch1 activation skews DP selection toward the CD8 SP lineage. Adult *CD4-CreERT2*; *R26YFP and CD4-CreERT2*; *R26ICN1-GFP* mice were subjected to the extended tamoxifen treatment regimen. (A) Schematic representation of the *ICN1-GFP^Ind^* and *YFP^Ind^* mouse models. (B) Flow cytometric analysis of thymocytes from 6-11 week-old *ICN1-GFP^Ind^* and *YFP^Ind^* mice. Representative sequential gating strategy for recently signalled, labelled thymocytes. *Left*: TCRβ^hi^ CD69^+^ cells among live single CD11b^-^ CD25^-^ thymocytes in each strain. *Middle*: YFP^+^ or GFP^+^ labelled cells among TCRβ^hi^ CD69^+^ cells in each strain. *Right*: CD4^-^ CD8α^+^ cells among labelled TCRβ^hi^ CD69^+^ cells in each strain. (C) Data were pooled from 2 experiments with n=5 per genotype.Scatter dot plots show the percentage of labelled cells among TCRβ^hi^ CD69^+^ thymocytes in each strain (*left*) or CD8 SP cells among labelled TCRβ^hi^ CD69^+^ thymocytes in each strain (*right*). Unpaired 2-tailed *t* test: *, *p*=0.03; ***, *p*<0.001.

DP thymocytes have an average lifespan of 3-4 days^54,55^, so we injected tamoxifen for 5 consecutive days in order to label 1-2 generations of DP thymocytes . SP thymocytes emigrate to the periphery within 4-5 days of being generated^56^. Therefore, we harvested thymocytes 3 days after the last tamoxifen injection, a time when most CD8 SP thymocytes derived from GFP^+^ or YFP^+^ DP thymocytes will still reside in the thymus. Because we were most interested in thymocytes undergoing selection, we evaluated tamoxifen-induced labelling among TCRβ^hi^ CD69^+^ cells in both strains (Fig. 3B). A similar percentage of TCRβ^hi^ CD69^+^ cells thymocytes were labelled in each strain, respectively, and labelled cells were mostly CD24^+^, indicating immaturity (Fig. 3B). Among labelled (GFP^+^ or YFP^+^) TCRβ^hi^ CD69^+^ “signaled” thymocytes, *ICN1-GFP^Ind^* mice had significantly more CD8 SP cells than *YFP^Ind^* mice (Fig. 3B). Collectively, these data that, like constitutive Notch1 activation, inducible Notch1 activation skews DP selection towards the CD8 SP fate.

### Notch1 activation prolongs DP survival and alters CD4/CD8 lineage determination during early CCR9^+^ stages of selection

A recent study showed that some single CD69^+^ DP thymocytes undergoing selection change express CD8 lineage markers such as *Cst7, Itgae* and *Nkg7*^57^, suggesting that these CD8 lineage characteristics are imprinted very early during selection. Interestingly, we found that although general maturation markers such as *Ccr7*, *Dntt, Rag1* and *Cd24a* were not differentially expressed by total DP thymocytes from *CD4-Cre* versus *ICN1-GFP^Ind^*mice, they expressed significantly higher levels of *Cst7, Itgae* and *Nkg7* than those from *CD4-Cre* mice (Suppl. Table 4). The CD8 lineage genes *Cd160, Xcl1* and *Gpr114* were also upregulated in DP thymocytes from *ICN1-GFP^Ind^*mice. These data suggest that Notch1 imposes a CD8 lineage bias during the earliest stages of selection.

To specifically test how Notch1 activation impacts early versus later stages of selection, we used a “time-stamp” tamoxifen pulse-chase strategy. *YFP^Ind^* and *ICN1-GFP^Ind^* mice were injected once with tamoxifen to label a single cohort of DP thymocytes. We gated on YFP^+^ or GFP^+^ cells and tracked their expression of CD4, CD8α, CD24, CD69, TCRβ, CCR9 and CCR7 after 2, 4 or 6 days of chase (Suppl. Fig. 1). After 2 days of chase, the frequency of DP cells among labelled thymocytes was similar in *YFP^Ind^* and *ICN1-GFI^Ind^*mice (Fig. 4A). By day 4, DP frequency among labelled *YFP^Ind^* thymocytes dropped to <10%, in keeping with the known turnover kinetics of these cells^54,58^. In striking contrast, DP frequency remained high among labelled thymocytes in *ICN1-GFP^Ind^*mice after 4 days, and a higher fraction were un-signaled CD69^-^ CCR7^-^ cells (Fig. 4B). However, BrdU-labelling of DP thymocytes was not higher in *ICN1-GFP^Con^*mice (Fig. 4C), suggesting that ICN1 prolonged DP survival rather than inducing their proliferation, in keeping with a prior study^59^. Indeed, by day 6, labelled DP thymocytes had largely disappeared in both strains. Collectively, these data suggest that CD4-Cre-mediated induction of Notch1 signalling increases the lifespan of DP thymocytes by about 2 days.

**Figure 4:**
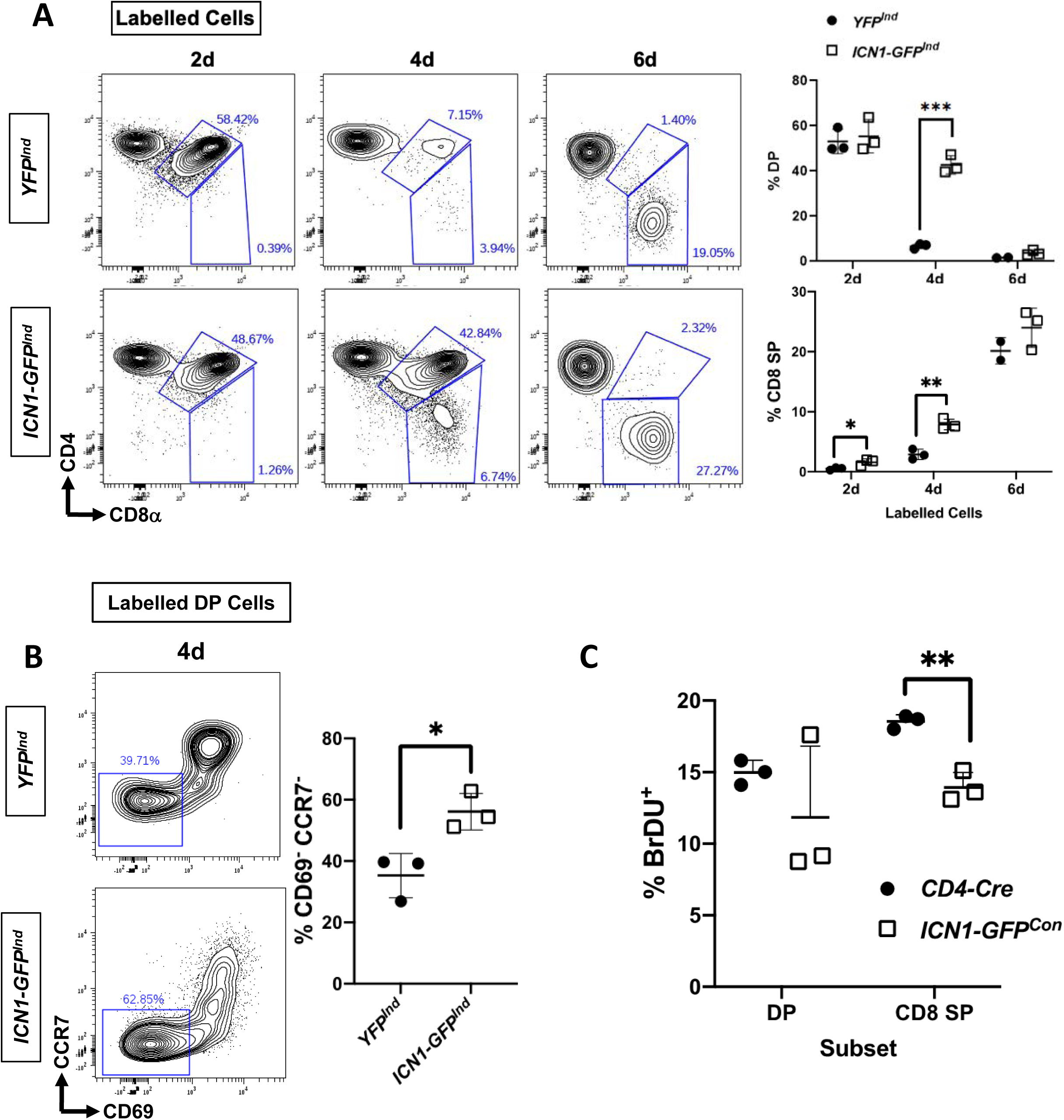
Activated Notch1 prolongs DP survival and skews selection toward the CD8 SP lineage. Time-stamp analysis of DP selection after tamoxifen-induced YFP or ICN1-GFP expression using the regimen shown in Suppl. Fig. 1. (A) Representative contour plots show CD4 versus CD8α on labelled (YFP^+^ or ICN1-GFP^+^) live single thymocytes 2, 4 and 6 days after tamoxifen injection. Scatter dot plots show the frequency of DP (*top*) and CD8 SP (*bottom*) cells among labelled thymocytes in each strain at each timepoint. Multiple t-tests, %DP: 4d, ***, q<0.001; %CD8 SP: 2d: *, q= 0.02; 4d: **, q=0.002. (B) Contour plots show CCR7 versus CD69 on labelled thymocytes after 2 and 4 days. Scatter dot plot shows the percentage of CD69^-^ CCR7^-^ among labelled DP thymocytes in each strain after 4 days. Unpaired 2-tailed t-test: *, p=0.02. Data is from 1 experiment with n=3 per genotype/per timepoint, except that n=2 for *YFP^Ind^* after 6 days. (C) Scatter dot plot shows the %BrDU^+^ cells among DP or CD8SP thymocytes as determined by flow cytometry 4h after injection of BrdU into *ICN-GFP^Con^* and *CD4-Cre* littermates. Data are shown from one experiment out of 2 performed.

By 2 days after tamoxifen administration, there were significantly more CD8 SP cells in *ICN1-GFP^Ind^* mice, and this difference increased by 4 days after tamoxifen injection (Fig. 4A), suggesting that the Notch acts early to bias selection towards the CD8 lineage. To examine how Notch1 impacts early versus later events during selection, we analyzed labelled TCRβ^hi^ CD69^+^ thymocytes at Stage 1 (CCR9^+^ CCR7^-^) versus Stage 2 (CCR9^+^ CCR7^+^) of selection. Stage 1 thymocytes were significantly more abundant among labelled signalled thymocytes in *ICN1-GFP^Ind^* mice after 4 days (Fig. 5A), which likely reflected the longer DP lifespan in this strain (Fig. 4). Interestingly, *ICN1-GFP^Ind^*mice had significantly fewer CD4^+^ CD8^lo^ cells among labelled signalled Stage 1 thymocytes (Fig. 5B, *left*). In addition, DP thymocytes were more abundant among labelled signalled Stage 2 thymocytes in *ICN1-GFP^Ind^* mice at both 2 and 4 days (Fig. 5B, *right*), again in keeping with the prolonged DP lifespan afforded by ICN1. Strikingly, there were significantly more CD8 SP thymocytes among labelled signalled Stage 2 cells in *ICN1-GFP^Ind^* mice, most notably after 4 days (Fig. 5B, *right*). Collectively, these experiments demonstrate that Notch1 activation at the DP stage impairs the emergence of CD4^+^ CD8^lo^ cells at Stage 1 and induces prominent CD8 lineage skewing by Stage 2 of thymocyte selection.

**Figure 5:**
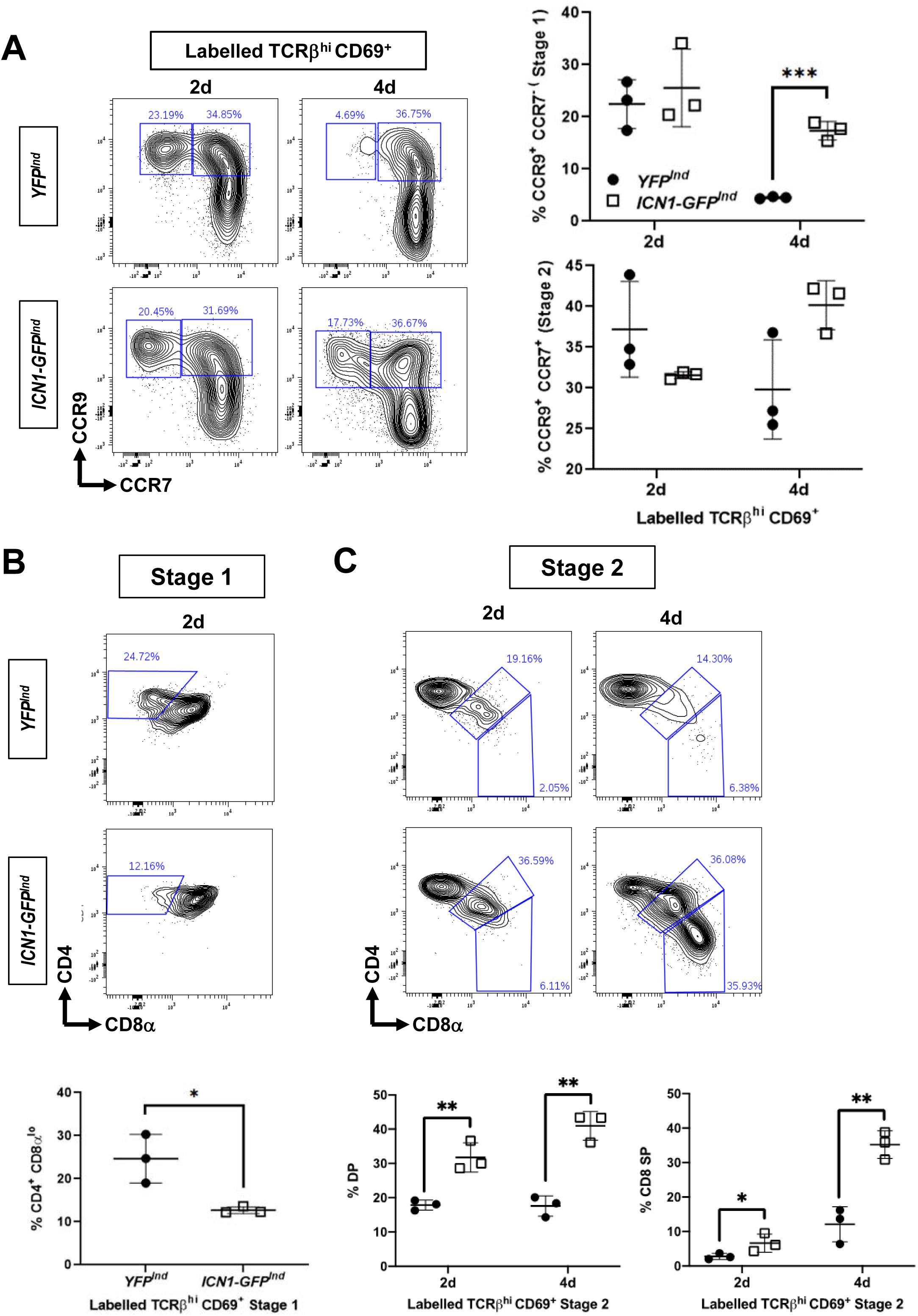
Activated Notch 1 alters CD4/CD8 lineage determination during early CCR9^+^ stages of selection. Data from the experiments shown in Fig. 4 were further analyzed to examine specific stages of selection. (A) Contour plots show CCR9 versus CCR7 on labelled TCRβ^hi^ CD69^+^ thymocytes from the indicated genotypes and timepoints. Scatter dot plots show the percentage of CCR9^+^ CCR7^-^ Stage 1 (*top*) and CCR9^+^ CCR7^+^ Stage 2 (*bottom*) among labelled TCRβ^hi^ CD69^+^ thymocytes in each strain. Multiple t-tests, ***, *q*<0.001. (B) Contour plots show CD4 versus CD8α on Stage 1 thymocytes at d2. Scatter dot plots show the frequency CD4^+^ CD8α^lo^ thymocytes. Unpaired 2-tailed t-test: *, *p*=0.02 (C) Contour plots show CD4 versus CD8α on Stage 2 thymocytes at d2 and d4. Scatter dot plots show the frequency DP or CD8 SP at each timepoint. Multiple t-tests, %DP: 2d, **, *q*=0.006; 4d: **, *q*=0.003. % CD8 SP: 2d, *, *q*=0.04; 4d: **, *q*=0.004.

### Notch1 activation blunts Gata3 up-regulation during Stage 1 of selection

To investigate how Notch1 activation skewed DP selection towards the CD8 SP fate at the expense of the CD4 SP fate, we evaluated expression of Gata3, ThPOK and Runx3, the key molecular determinants CD4/CD8 lineage determination, in *CD4-Cre* versus *ICN1-GFP^Con^* mice. Signalled DP thymocytes had similarly low frequencies of Runx3^+^ cells in both strains (Suppl. Fig. 2), suggesting that skewing towards the CD8 SP fate does not result from premature Runx3 induction in *ICN1-GFP^Con^*DP thymocytes.

We next asked whether Notch1 activation compromised upregulation of Gata3 or ThPOK during selection since prior studies have shown that loss-of-function mutations in either gene prevents CD4 commitment and allows redirection of pMHCII-restricted DP thymocytes to the CD8 lineage^31,32^. Gata3 acts upstream of ThPOK^26^, so we examined the expression of these proteins in signalled thymocytes at Stage 1 and Stage 2 of selection. First, we confirmed that the phenotypic “time-stamp” method in the *ICN1-GFP^Con^*model recapitulated our findings using physical time-stamping in the *ICN1-GFP^Ind^*model. Namely, *ICN1-GFP^Con^* mice had fewer CD4^+^ CD8^lo^ cells during Stage 1 and more CD8 SP cells in Stage 2 (Fig. 6A). Notably, there were also significantly fewer Gata3^hi^ CD8^lo^ cells among signalled Stage 1 thymocytes from *ICN1-GFP^Ind^* mice, and very few had emerged by Stage 2 (Fig. 6B). In addition, CD4^+^ ThPOK^+^ cells were prominent in signalled Stage 2 thymocytes from *CD4-Cre* but not *ICN1-GFP^Con^* mice (Fig. 6C). These findings suggest that activated Notch1 blunts Gata3 up-regulation and inhibits the emergence of CD4^+^ CD8^lo^ selection intermediates during Stage 1, thereby preventing ThPOK induction and CD4 lineage emergence in response to pMHCII in Stage 2.

**Figure 6:**
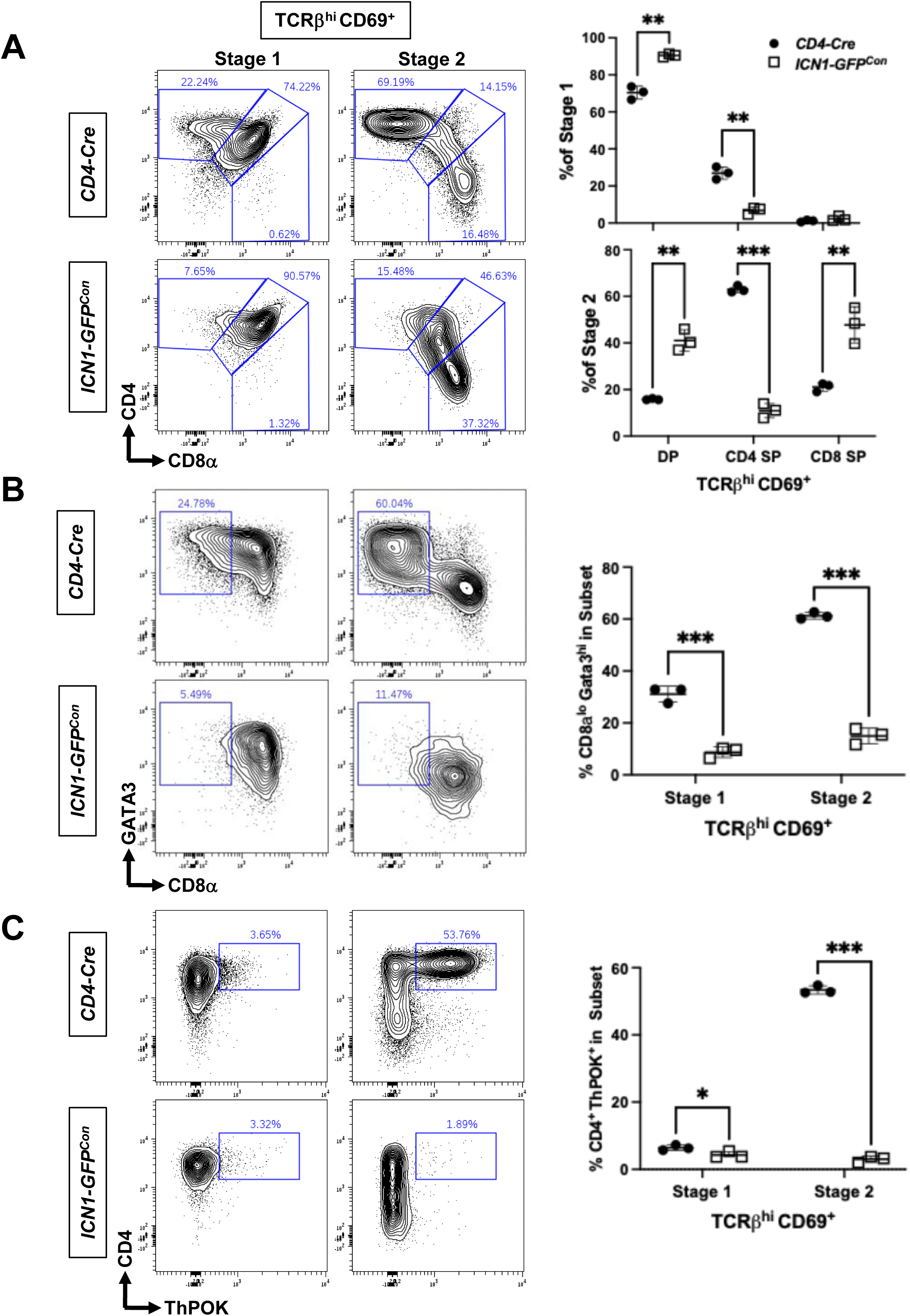
Activated Notch 1 blunts Gata3 upregulation and prevents ThPOK induction. Flow cytometric analysis of Gata3 and ThPOK expression during selection stages at steady state in each strain. (A) Contour plots showing CD4 versus CD8 on TCRβ^hi^ CD69^+^ cells during Stage 1 (CCR9^+^ CCR7^-^) and Stage 2 (CCR9^+^ CCR7^+^) of selection. Scatter dot plots show the frequency of DP, CD4 SP and CD8 SP thymocytes within Stage 1 and Stage 2 of selection in each strain. Multiple t-tests, **, *q*=0.002. (B) Contour plots showing Gata3 versus CD8 on TCRβ^hi^ CD69^+^ Stage 1 and Stage 2 of selection. Scatter dot plot shows the frequency of Gata3^hi^ CD8^lo^ thymocytes within Stage 1 and Stage 2 of selection in *ICN1-GFP^Con^* mice and littermate controls. Multiple t-tests, ***, *q*<0.001. (C) Contour plots showing CD4 versus ThPOK on TCRβ^hi^ CD69^+^ Stage 1 and Stage 2 of selection. Scatter dot plot shows the frequency of CD4^+^ ThPOK^+^ thymocytes within Stage 1 and Stage 2 of selection in each strain. Data is from 3 experiments with n=3 per genotype/per timepoint. Multiple t-tests, *, *q*=0.04; ***, *q*<0.001.

### Notch1 activation re-directs pMHCII-specific DP thymocytes to the CD8 lineage

CD4^+^ CD8^lo^ thymocytes are thought to serve as intermediates not only for selection of CD4 SP cells by pMHCII, but also for selection of CD8 SP cells signalled by pMHCI. It was thus surprising that activated Notch1 skewed DP selection towards the CD8 lineage while also impairing generation of CD4^+^ CD8^lo^ selection intermediates. Therefore, we used the OT-I transgenic^60^ TCRαβ model to ask how activated Notch1 impacted generation of CD4^+^ CD8^lo^ cells specifically in response to pMHCI signalling. As previously reported, OT-I thymocytes have abundant CD4^+^ CD8^lo^ selection intermediates, but few CD4 SP thymocytes^60^ (Fig. 7A). However, activated Notch1 strongly impaired generation of CD4^+^ CD8^lo^ thymocytes. Activated Notch1 also increased production of CD8 SP cells in *ICN1-GFP^Con^*; *OT-I* mice, suggesting that it improved OT-I selection efficiency. These data show that activated Notch1 diverts pMHCI signalled DP thymocytes to the CD8 SP stage without generating CD4^+^ CD8^lo^ intermediates.

**Figure 7:**
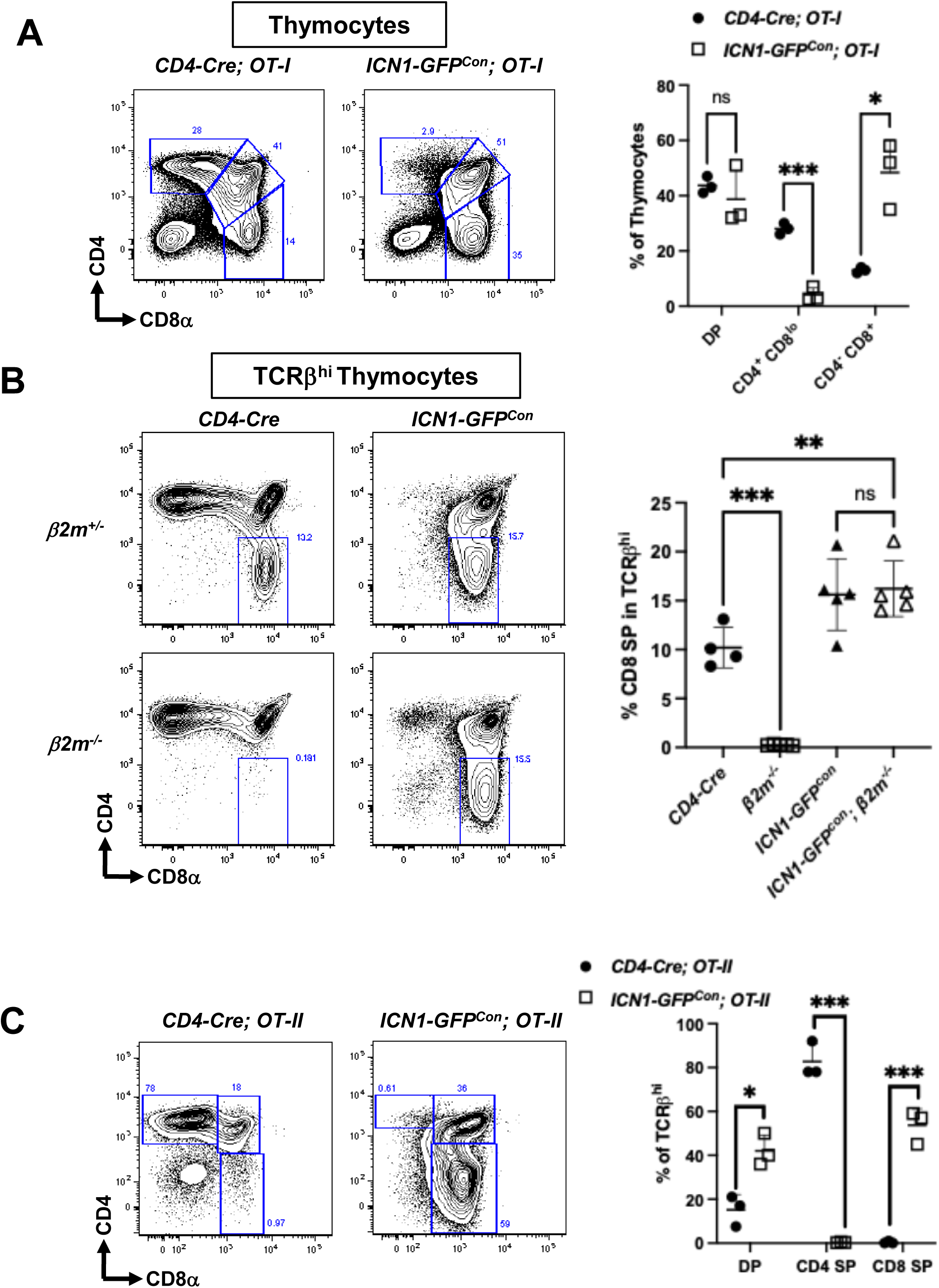
Impact of activated Notch1 on selection of pMHCI- and pMHCII-specific DP thymocytes. Flow cytometric analysis of CD4^+^CD8^lo^, CD4 and CD8 SP thymocytes in *ICN1-GFP^Con^* mice expressing transgenic pMHCI- or pMHCII-specific TCRαβ or β*2m*^-/-^ mice lacking MHCI. Contour plots show CD4 versus CD8 on total or TCRβ^hi^ thymocytes in each strain. (A) Scatter dot plots show the frequency of DP, CD4^+^ CD8^lo^, and CD8 SP cells. Multiple *t* tests, *, *q*=0.01; ***, *q*<0.001. (B) Scatter dot plots show the frequency of CD8 SP cells in each strain. One-way ANOVA with post-hoc *t* tests comparing each strain to *ICN1-GFP^con^*, **, *q*=0.006; ***, *q*<0.001 (C) (Scatter dot plots show the frequency of DP, CD4 SP and CD8 SP cells in each strain. Multiple t-tests, *, *q*=0.01; ***, *q*<0.001.

The ability of Notch1 activation in DP thymocytes to blunt Gata3 induction during selection suggests that skewing towards the CD8 SP lineage in *ICN1-GFP^Con^*mice could result from re-direction of pMHCII-specific DP thymocytes to the CD8 lineage. To test this notion, we generated *ICN1-GFP^Con^*; β*2m^-/-^* mice, which lack MHCI. As expected, *CD4-Cre*; β*2m^-/-^* mice developed only CD4 SP thymocytes (Fig. 7B). However, *ICN1-GFP^Con^;*β*2m^-/-^*mice had similar frequencies of TCRβ^hi^ CD8 SP thymocytes as *ICN1-GFP^Con^*mice, and significantly more than *CD4-Cre* mice. These findings demonstrate that ICN1 robustly promotes development of CD8 SP thymocytes in the absence of MHCI, likely in response to pMHCII.

To formally test the idea that ICN1 can re-direct pMHCII-specific DP thymocytes to the CD8 lineage, we generated *ICN1-GFP^Con^ Rag2^-/-^* mice expressing the OT-II transgenic TCRαβ which recognizes a pMHCII complex expressed by B6 thymic epithelial cells^61^. As expected, most TCRβ^hi^ cells in *CD4-Cre; OT-II* mice were CD4 SP (Fig. 7C). *ICN1-GFP^Con^* increased the frequency of DP thymocytes among TCRβ^hi^ thymocytes, as we observed in polyclonal *ICN1-GFP^Ind^* mice. More notably, ICN1 robustly blocked pMHCII-specific DP from generating CD4 SP in *ICN1-GFP^Con^*; *OT-II Rag2^-/-^* mice and diverted them to the CD8 lineage, demonstrating that ICN1 could re-direct DP thymocytes expressing a monoclonal pMHCII-specific TCRαβ into the CD8 lineage.

## DISCUSSION

Notch1 signalling critically regulates early T cell development but, its role in positive selection and CD4/CD8 lineage determination of DP thymocytes has remained elusive, in part because early studies expressing activated Notch1 in earlier T cell progenitors had inconsistent findings. Here, we re-addressed this question by using CD4-Cre to activate Notch1 specifically in DP thymocytes, which circumvented the induction of thymic lymphomas. In accordance with previous findings using *LckPr-ICN1* mice^13,19^, we showed that CD4-Cre-mediated induction of canonical, *Rbpj*-dependent Notch1 signalling potently inhibits CD4 lineage determination and skews DP selection to the CD8 lineage. Using a tamoxifen-inducible CD4-CreER^T2^ “time-stamp” model, we showed that Notch1 acts during the earliest stage of DP selection to impair generation of CD4^+^ CD8^lo^ selection intermediates and to promote the CD8 fate at the expense of the CD4 fate. Finally, we showed that ICN1 blunts Gata3 upregulation in early selection intermediates, impairing ThPOK induction and allowing re-direction of pMHCII-specific DP precursors to the CD8 lineage. Collectively, these demonstrate that Notch1 activation in DP thymocytes acts on early selection intermediates to block Gata3-mediated differentiation of CD4-fated, pMHCII-specific DP precursors and re-direct them to the CD8 lineage.

In contrast to studies that used ectopic activation of Notch1 in HSC or DN2/DN3 thymocytes, we found that CD4-Cre-induced Notch1 activation did not induce *Myc*, increase DP thymocyte proliferation or cause T cell leukaemia/thymic lymphoma. Nonetheless, time-stamp studies showed that activated Notch1 slightly increased the lifespan of both pre- and post-selection DP thymocytes, in accordance with a study showing that ICN1 up-regulates Bcl2 in thymocytes^59^. Interestingly, Korsemeyer and colleagues^62^ showed that transgenic *hBCL2* increased the frequency of CD8 SP thymocytes, suggesting that improving thymocyte survival has a lineage-specific impact on positive selection. However, we found that although the same *hBCL2* transgene significantly improved DP thymocyte viability in *ICN1-GFP^Con^* mice, it did not restore their generation of TCRβ^hi^ CD4 SP thymocytes. Therefore, ICN1-induced Bcl2 did not account for the selective impact of Notch1 activation on CD4/CD8 lineage determination in our study.

We used a tamoxifen-inducible time-stamp system to study the developmental dynamics of Notch1 activation on CD4/CD8 lineage determination. The kinetics of Runx3 up-regulation were not perturbed by Notch1 activation, suggesting that premature Runx3 expression in DP thymocytes did not cause CD8 lineage skewing in our models. However, Notch1 activation blunted Gata3 up-regulation and reduced the generation of CD4^+^ CD8^lo^ cells in signalled CCR7^-^ Stage 1 thymocytes. Gata3 up-regulation controls a pre-commitment checkpoint for the CD4 lineage, independently of its ThPOK-inducing function^26^, so it is likely that the reduction in Gata3 is mechanistically linked to the compromised generation of CD4 cells in our *ICN1-GFP^Con^*and *ICN1-GFP^Ind^* models. In support of this notion, generation of CD4^+^ CD8^lo^ cells, which occurs in response to DP signalling by pMHCI or pMHCII^63,64^, was greatly reduced by Notch1 activation, similar to the impact of Gata3 deficiency in DP thymocytes^26^. Thus, Notch1 activation and Gata3 loss have similar impacts on the early stages of DP selection, and both manipulations largely prevent the emergence of CD4^+^ ThPOK^+^ cells among CCR7^+^ cells at Stage 2. However, in contrast to the blunting of Gata3 up-regulated by Notch1 activation in DP thymocytes, Notch activation directly up-regulates Gata3 during early T cell development^65,66^ and in Th2 polarization^67^. Thus, Notch1 can mediate opposite effects on Gata3 expression in different developmental contexts, likely due to differences in the transcriptional and/or epigenetic landscapes of the cells at each stage of differentiation.

Previous studies of *LckPr-ICN1* mice showed that Notch1 activation could restore selection of CD8 SP to β*2m^-/-^* mice lacking MHCI^13^. We confirmed this finding for CD4-Cre-induced Notch1 activation in *ICN1-GFP^Con^*; β*2m^-/-^*mice. However, these mice express a diverse, polyclonal TCRαβ repertoire, so it was unclear whether Notch1 activation could re-direct a pMHCII-specific TCRαβ towards the CD8 fate. We resolved this question by showing that DP thymocytes expressing the pMHCII-specific OT-II TCRαβ were re-directed to the CD8 SP fate with high efficiency. These findings provide direct evidence that Notch1 activation can over-ride mechanisms that match TCRαβ specificity for MHC to CD4/CD8 co-receptor expression to skew DP selection towards the CD8 lineage.

In conclusion, our study used constitutive and inducible genetic models combined with high dimensional flow cytometry to provide the first kinetic analysis of the effect of activated Notch1 on selection and CD4/CD8 lineage determination. These studies resolve the long-standing uncertainty about how activated Notch1 impacts DP thymocyte selection and provide novel insights into mechanisms by which activated Notch1 impacts molecular determinants of the CD4/CD8 lineage decision.

## Supporting information

supplemental figures with legend

Suppl. Tables Illumina Microarray

## ACKNOWLEDGEMENTS

We acknowledge awards from the Vanier Canada Graduate Scholarship (ME); the SickKids Hospital Research Training Center (Restracomp awards to ME and MSG); and the University of Toronto Department of Immunology Doctoral Completion Award (ME). This work was also supported by operating grants to CJG from the Natural Sciences and Engineering Research Council of Canada (NSERC, funding reference number 4018370-12), the Canadian Institutes of Health Research (CIHR, funding reference numbers 11530, 165973) and Genome Canada through the Ontario Genomics Institute. We thank Dr. Shaheena Bashir for analyzing the microarray data and Dr. Juan Carlos Zuniga Pflucker (JCZP) of the Toronto Sunnybrook Research Institute, Toronto for providing B6.*Rbpj^f^* mice. Key aspects of this work were supported by the Flow Cytometry Facility and the Center for Applied Genomics research core facilities at the Hospital for SickKids Research Institute.

## AUTHOR CONTRIBUTIONS

CJG designed the study with input from PCK, AB and ME, who together with the other co-authors also performed experiments and analyzed data. ME and CJG wrote the manuscript.

## METHODS

### Mice

Mice were obtained from the sources shown in Table 1 below. The *CD4-Cre* and *R26^ICN1-GFP^*strains were each backcrossed 10 generations to C57BL/B6J (B6). The resulting *B6.CD4-Cre* and *B6.R26^ICN1-GFP^* strains were then intercrossed to generate *ICN-GFP^Con^* mice that express sequences encoding the intracellular Notch1 (ICN1) region (amino acids 1749-2293), an internal ribosomal entry sequence (IRES) flanked by loxP sites, and GFP inserted into the *R26* locus in DP thymocytes and their progeny. *ICN-GFP^Con^* mice and their *CD4-Cre^-^* or *R26^ICN1-GFP^*^-^ littermates used in this study were 3-6 weeks old. *CD4CreER^T2^* mice were bred to either *R26^ICN1-GFP^* or *R26^EYFP^* mice to obtain *CD4-CreER^T2^; R26^ICN1-GFP^* (*ICN-GFP^Ind^*) and *CD4-CreER^T2^; R26^EYFP^* (*YFP^Ind^*) mice, respectively. 4-15-week-old mice were used for all experiments. *ICN-GFP^Con^* mice were intercrossed with *OT-II* or β*2m^-/-^* mice for multiple generations to generate *ICN-GFP^Con^; OT-II* and *ICN-GFP^Con^;* β*2m^-/-^* mice, respectively and their *CD4-Cre* littermate controls.

**Table 1:** Mouse strain information.

| Official Name | Source | Strain ID | Short name |
| --- | --- | --- | --- |
| Tg(Cd4-cre)1Cwi/BfluJ | JAX | JAX 017336 | *CD4-Cre* |
| B6(129X1)-Tg(Cd4-cre/ERT2)11Gnri/J | JAX | JAX 022356 | *CD4-CreER<sup>T2</sup>* |
| B6.129X1-Gt(ROSA)26Sor<sup>tm1(EYFP)Cos</sup>/J | JAX | JAX 006148 | *R26<sup>EYFP</sup>* |
| B6.Cg-Tg(TcraTcrb)425Cbn/J | JAX | JAX 004194 | *OT-II* |
| C57BL/6-Tg(TcraTcrb)1100Mjb/J | JAX | JAX 003831 | *OT-I* |
| B6.129P2-*B2m<sup>tm1Unc</sup>*/DcrJ | JAX | JAX 002087 | *B2m KO* |
| Gt(ROSA)26Sor<sup>tm1(NOTCH1)Dam</sup>/J | JAX | JAX 008159 | *R26<sup>ICN1-GFP</sup>* |
| C3H/He-Tg(LCKprBCL2)36Sjk/J | JAX | JAX 002427 | *hBCL2-Tg* |
| B6.Cg-Rbpj<sup>tm1Hon</sup> | JCZP | MGI 3583755 | *Rbpj<sup>f</sup>* |
| C57BL/6J | JAX | JAX 000664 | *B6* |

All animals were maintained and bred in specific pathogen-free conditions in the animal facilities of SickKids Peter Giligan Center for Research and Learning (PGCRL) and the Centre for Phenogenomics (TCP) and euthanized by CO_2_ or cervical dislocation. All animal work was approved by the Institutional Animal Care Committees. Mouse genotypes were determined by polymerase chain reaction (PCR) amplification of tail deoxyribonucleic acid (DNA). Tail DNA was isolated using the DNeasy Tissue Kit (Qiagen) according to the manufacturer’s instructions, Primers and PCR conditions used to genotype each strain are available on request.

### Tamoxifen studies and time stamp labelling

For the extended treatment studies, *ICN1-GFP^ind^* and *YFP^ind^* mice were injected intraperitoneally for 5 consecutive days with 2 mg tamoxifen (Sigma-Aldrich, Catalogue number T5648-1G) emulsified in sunflower oil. Mice were sacrificed 3 days after the last injection. For the time-course experiments, mice were injected intraperitoneally with 2 mg tamoxifen on day 0 and sacrificed 2, 4 or 6 days later.

### Thymus isolation

Thymii were harvested, chopped finely with scissors and crushed with the plunger of a 3 ml syringe onto a 100 µm wire mesh into a petri dish. The metal mesh was rinsed with 5-10 ml staining media (SM) consisting of Hank’s Balanced Salt Solution (HBSS, Wisent Inc.) supplemented with 10 mM HEPES pH 7.2 (Wisent Inc.) and 5% calf serum (Wisent Inc.) 2-3 times to collect single-cell suspension. The single-cell suspension was passed through a 70 µm filter (BD Biosciences) on a 50 ml falcon tube and the filter was then washed with another 10 ml SM. Cells were then pelleted at 300-400 *x g*, resuspended in 10-20 ml of SM and viable cell counts were determined by Trypan Blue staining using a hemocytometer.

### Flow Cytometry

The fluorochrome-conjugated antibodies (Ab) used in this study are shown in Table 2. Ab cocktails were prepared by diluting each Ab in SM to saturating concentrations that were pre-determined by titration. Single-cell suspensions of thymocytes (2 x10^6^) cells were washed in 1 ml SM and then re-suspended in 50 µl of Ab cocktail for 30’ at room temperature (RT) in the dark. To stain chemokine receptors, cells were pre-incubated with anti-chemokine receptor Abs in 25 µl at 2X final concentration for 10’ at 37 °C. The remaining anti-surface marker Abs were then added in 25 µl at 2X final concentration and cells were shifted to RT for an additional 20’. After staining, cells were washed by adding 0.5 ml of SM followed by pelleting 300xg for 5’. For Ab panels that included a biotinylated Ab, cells were then stained in 50 µl of fluorochrome-conjugated Streptavidin (Av) for 30’ at RT in the dark. Following antibody staining, cells were washed with 1 ml serum-free PBS prior to staining with pre-optimized concentrations of Zombie UV or FVS-510 fixable viability dye in PBS for 30’ at room temperature. Cells were then washed with 0.5 ml SM. In some cases, cells were subsequently fixed, permeabilized and stained with Abs specific for transcription factors following the manufacturer’s protocol supplied with the BD Transcription Factor Buffer Set. For all post-fixation steps, cells were pelleted at 800xg for 5’. After completing all staining steps, cells were resuspended in 0.3-0.5 ml of SM and filtered through a 40 or 70 µm nylon mesh immediately before analysis.

**Table 2:**
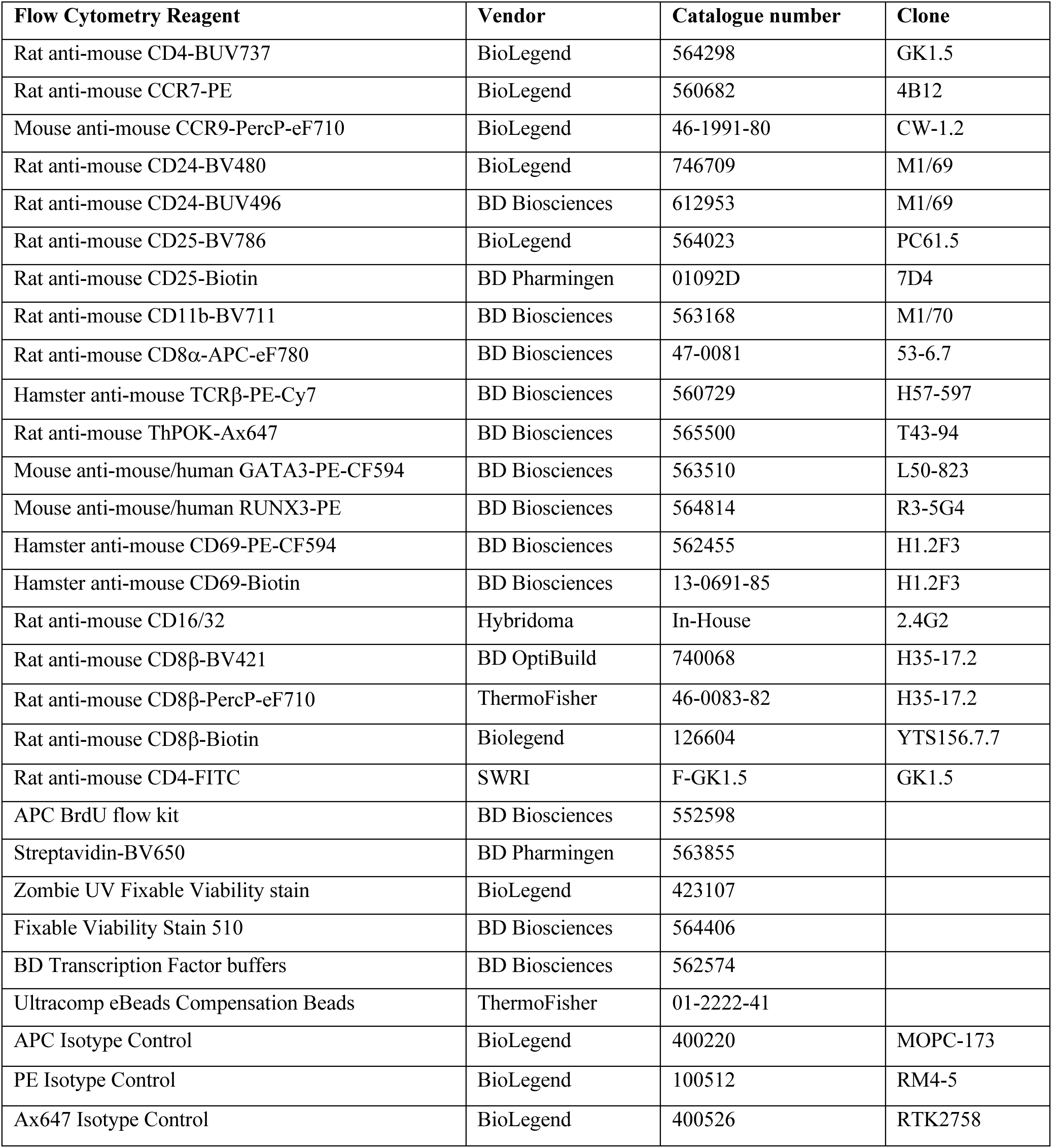
Flow cytometry reagents.

| Flow Cytometry Reagent | Vendor | Catalogue number | Clone |
| --- | --- | --- | --- |
| Rat anti-mouse CD4-BUV737 | BioLegend | 564298 | GK1.5 |
| Rat anti-mouse CCR7-PE | BioLegend | 560682 | 4B12 |
| Mouse anti-mouse CCR9-PercP-eF710 | BioLegend | 46-1991-80 | CW-1.2 |
| Rat anti-mouse CD24-BV480 | BioLegend | 746709 | M1/69 |
| Rat anti-mouse CD24-BUV496 | BD Biosciences | 612953 | M1/69 |
| Rat anti-mouse CD25-BV786 | BioLegend | 564023 | PC61.5 |
| Rat anti-mouse CD25-Biotin | BD Pharmingen | 01092D | 7D4 |
| Rat anti-mouse CD11b-BV711 | BD Biosciences | 563168 | M1/70 |
| Rat anti-mouse CD8α-APC-eF780 | BD Biosciences | 47-0081 | 53-6.7 |
| Hamster anti-mouse TCRβ-PE-Cy7 | BD Biosciences | 560729 | H57-597 |
| Rat anti-mouse ThPOK-Ax647 | BD Biosciences | 565500 | T43-94 |
| Mouse anti-mouse/human GATA3-PE-CF594 | BD Biosciences | 563510 | L50-823 |
| Mouse anti-mouse/human RUNX3-PE | BD Biosciences | 564814 | R3-5G4 |
| Hamster anti-mouse CD69-PE-CF594 | BD Biosciences | 562455 | H1.2F3 |
| Hamster anti-mouse CD69-Biotin | BD Biosciences | 13-0691-85 | H1.2F3 |
| Rat anti-mouse CD16/32 | Hybridoma | In-House | 2.4G2 |
| Rat anti-mouse CD8β-BV421 | BD OptiBuild | 740068 | H35-17.2 |
| Rat anti-mouse CD8β-PercP-eF710 | ThermoFisher | 46-0083-82 | H35-17.2 |
| Rat anti-mouse CD8β-Biotin | Biolegend | 126604 | YTS156.7.7 |
| Rat anti-mouse CD4-FITC | SWRI | F-GK1.5 | GK1.5 |
| APC BrdU flow kit | BD Biosciences | 552598 |  |
| Streptavidin-BV650 | BD Pharmingen | 563855 |  |
| Zombie UV Fixable Viability stain | BioLegend | 423107 |  |
| Fixable Viability Stain 510 | BD Biosciences | 564406 |  |
| BD Transcription Factor buffers | BD Biosciences | 562574 |  |
| Ultracomp eBeads Compensation Beads | ThermoFisher | 01-2222-41 |  |
| APC Isotype Control | BioLegend | 400220 | MOPC-173 |
| PE Isotype Control | BioLegend | 100512 | RM4-5 |
| Ax647 Isotype Control | BioLegend | 400526 | RTK2758 |

For compensation controls, isotype controls were titrated on Ultracomp compensation beads (ThermoFisher Scientific) to determine the dilution for the highest, on-scale MFI to be used for compensation non-tandem dye-conjugated Abs. For tandem dye-conjugated Abs, all Abs were titrated on Ultracomp compensation beads to determine the dilution for the highest, on-scale MFI. For each experiment, 25ul of Ultracomp compensation beads were stained with 25ul of either isotype controls, for non-tandem dye-conjugated Abs, or the same Abs used to stain the cells, for tandem dye-conjugated Abs, for 30’minutes at 2X the pre-determined optimal dilution. If cells underwent fixation, all compensation controls were fixed for the same length of time, under the same conditions.

Fluorescence was analyzed on a BD LSR-Fortessa running FACSDiva v8.0.1. Instrument settings were pre-optimized and standardized using single-stained compensation control beads and unstained thymocytes using the principles established by Maecker and Trotter^68^. Briefly, the BD Cytometer Set-up and Tracking beads and FACSDiva v8.0.1 software was used according to the manufacturer’s instructions to define and perform daily QC monitoring of baseline performance metrics for each detector used in the instrument configuration used in this study. We then defined a “minimal noise” target and corresponding target Median Fluorescence Intensity (MFI) values for each channel on Rainbow calibration particles, 4^th^ peak from the 8-peak set (Spherotech, RCP-30-5A-4) that were recorded prior to performing experiments. Set up of the cytometer on the day of the experiment immediately before sample acquisition was done by running Rainbow calibration particles, 4^th^ peak from the 8 peak set and determining the voltage required to attain the pre-determined target MFI value for each channel. For IEL samples, to determine absolute cell numbers, each sample was spiked with CountBright^TM^ Absolute Counting Beads (ThermoFisher, C36950) according to manufacturer’s protocol. Flow data were analyzed using FlowJo v10 and Cytobank and pre-gating to exclude debris, dead cells and doublets. Additional pre-gates for each plot are stated for each figure.

### Gene-expression profiling by Illumina microarrays

The RNeasy isolation kit (Qiagen) was used to isolate total RNA from sorted DP and CD8 SP thymocytes sorted from 4–6-week-old *ICN1-GFP^Con^* mice and *WT* littermates (n=4/genotype). Illumina MouseRef-8 v2.0 Expression BeadChip kits were used for genome-wide expression profiling according to standard protocols at The Centre for Applied Genomics at the Hospital for Sick Children. We used *Limma* (linear models for microarray data) and the Benjamini-Hochberg method to estimate the false-discovery rate (FDR) in R Bioconductor 2.13.0 as previously described^69^. We thus identified 366 probe sets that were significantly differentially expressed between genotypes (FDR-adjusted *q* value *<*10^-4^) with a Log2 mean *ICN1-GFP^Con^/*mean *CD4-Cre* fold-change (FC) ratio of ≤ −1.0 (down-regulated genes) or ≥1.0 (up-regulated genes). This filtering process yielded 180 significantly up-regulated unique genes in *ICN1-GFP^Con^*CD8 SP thymocytes (Log2 FC ratio 1.0 to 6.41 and *q*≤10^-4^) and 119 that were significantly down-regulated (Log2 FC ratio −1.0 to −4.23 and *q*≤10^-4^). The MyGeneSet tool from the Immunological Genome Project (ImmGen) website^70^ was used to visualize the expression of selected down-regulated or up-regulated genes among immature and mature T cell subsets subjected to global gene expression profiling by the Immunological Genome Project Consortium (Microarray V1 data set).

### Statistical analyses

Prism 8.0 was used for statistical analysis and graphing. For comparison of two groups, the unpaired two-tailed Student’s *t-*test was used (α=0.05). For unpaired multiple *t*-tests and ANOVA (for comparison of 3 or more groups), the Holm-Sidak method was used to correct for multiple testing (α=0.05) yielding *q* values. Details on sample size, experimental replicates and statistics are included in the Figure Legends.

