## supplemental figures with legend for "Activated Notch1 Redirects CD4-Fated CD4^+^ CD8^+^ Precursors to the CD8 Lineage During Thymocyte Selection Without Causing T Cell Leukemia"

### Supplementary Figure 1

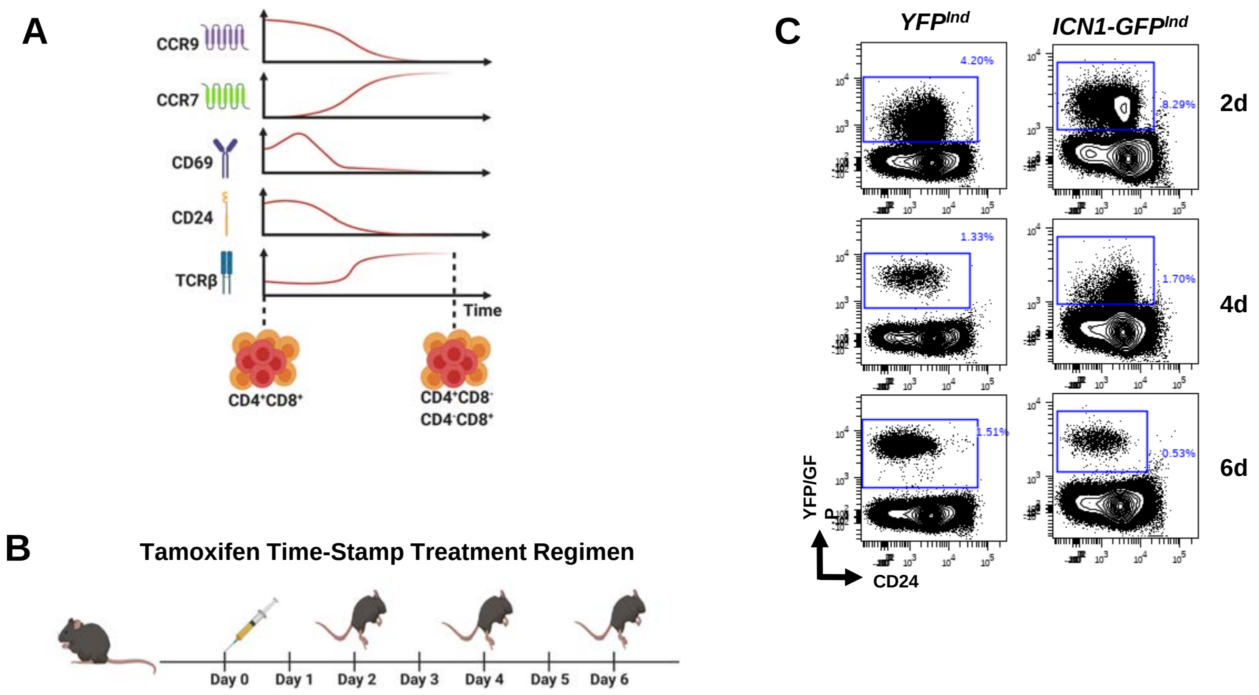

**Supplementary Figure 1: Time-stamp approach to follow DP thymocyte fate after tamoxifen-induced labelling in *YFP*$^{Ind}$ versus *ICN1-GFP*$^{Ind}$ mice**

(A) Schematic representation of changes in expression of CCR9, CCR7, CD69, CD24 and TCRβ DP thymocytes transit from pre-selection to fully mature CD4 and CD8 SP thymocytes.

(B) Time-stamp regimen used to follow DP thymocyte selection outcomes analyzed 2, 4 and 6 days after a single tamoxifen injection.

(C) YFP or GFP versus CD24 gating of CD11b$^-$ CD25$^-$ live singlets used to identify labelled thymocytes at each time-point in *YFP*$^{Ind}$ versus *ICN1-GFP*$^{Ind}$ mice in Figure 4.

### Supplementary Figure 2

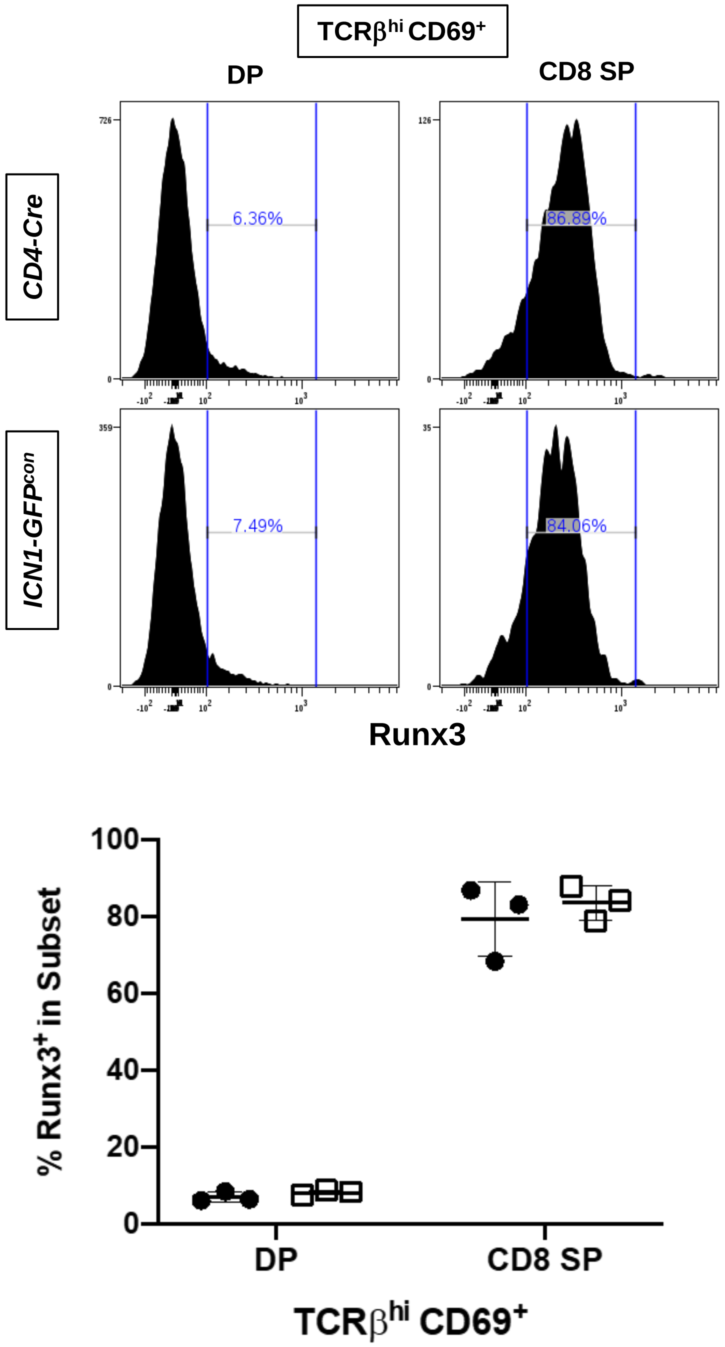

**Supplementary Figure 2: CD4-Cre-induced Notch1 activation does not induce early up-regulation of Runx3**

Histograms were gated to enumerate $Runx3^{+}$ cells among $TCR\beta^{hi}$ $CD69^{+}$ DP and CD8 SP thymocytes. Scatter dot plot shows the frequency of $Runx3^{+}$ thymocytes within each subset by strain. Data are shown from one experiment out of 2 performed.
