## Supplementary material for "Activated Notch1 Redirects CD4-Fated CD4^+^ CD8^+^ Precursors to the CD8 Lineage During Thymocyte Selection Without Causing T Cell Leukemia": Suppl. Tables Illumina Microarray

### Supplementary Table 1: Higher expression of selected direct Notch target genes in *ICN1-GFP*$^{Con}$ DP thymocytes

| Probe_Id | Symbol | Gene Name | Entrez Gene ID | Mean Expression in DP thymocytes (Log2) *CD4-Cre* | *ICN1-GFP*$^{Con}$ | Log2 FC | *FDR* |
|---|---|---|---|---|---|---|---|
| ***A. Global Direct Notch Targets*** | | | | | | | |
| ILMN_2641793 | *Dtx1* | *deltex 1 homolog (Drosophila)* | 14357 | 6.99 | 12.97 | 5.99 | 1.60E-20 |
| ILMN_1247691 | *Hes1* | *hairy and enhancer of split 1 (Drosophila)* | 15205 | 6.72 | 10.55 | 3.83 | 6.63E-18 |
| ILMN_2737416 | *Fjx1* | *four jointed box 1 (Drosophila)* | 14221 | 6.44 | 9.66 | 3.22 | 1.52E-10 |
| ILMN_2690596 | *Hey1* | *hairy/enhancer-of-split related with YRPW motif 1* | 15213 | 6.84 | 9.90 | 3.06 | 1.49E-10 |
| ILMN_1241915 | *Notch1* | *Notch gene homolog 1 (Drosophila)* | 18128 | 9.93 | 12.09 | 2.17 | 2.01E-10 |
| ILMN_1213139 | *Heyl* | *hairy/enhancer-of-split related with YRPW motif-like* | 56198 | 6.61 | 7.46 | 0.85 | 1.49E-10 |
| ILMN_2624153 | *Hes5* | *hairy and enhancer of split 5 (Drosophila)* | 15208 | 6.64 | 6.68 | 0.04 | 4.45E-03 |
| ***B. Direct Notch Targets in Immature T cells*** | | | | | | | |
| ILMN_1254670 | *Ptcra* | *pre T-cell antigen receptor alpha* | 19208 | 8.36 | 11.22 | 2.86 | 1.82E-12 |
| ILMN_2697380 | *Notch3* | *Notch gene homolog 3 (Drosophila)* | 18131 | 7.41 | 9.47 | 2.07 | 3.70E-07 |
| ILMN_2596979 | *Nrarp* | *Notch-regulated ankyrin repeat protein* | 67122 | 7.93 | 8.19 | 0.26 | 5.93E-09 |
| ILMN_2717975 | *Tcf7* | *transcription factor 7, T-cell specific* | 21414 | 11.82 | 11.74 | -0.08 | 3.09E-02 |
| ILMN_1233474 | *Il2ra* | *interleukin 2 receptor, alpha chain* | 16184 | 6.77 | 7.26 | 0.49 | 5.70E-02 |
| ILMN_2659739 | *Il7r* | *interleukin 7 receptor* | 16197 | 6.72 | 6.93 | 0.21 | 2.38E-01 |
| ***C. Direct Notch Targets in T Cell Leukemia*** | | | | | | | |
| ILMN_1231146 | *Agfg1* | *ArfGAP with FG repeats 1* | 15463 | 9.68 | 11.17 | 1.48 | 4.85E-10 |
| ILMN_3163001 | *Depdc6* | *DEP domain containing 6* | 97998 | 7.03 | 7.25 | 0.22 | 5.04E-05 |
| ILMN_2623526 | *Myc* | *myelocytomatosis oncogene* | 17869 | 6.96 | 7.03 | 0.08 | 5.75E-01 |
| ILMN_1239469 | *Lef1* | *lymphoid enhancer binding factor 1* | 16842 | 8.62 | 9.65 | 1.03 | 9.66E-01 |

*Legend:* The table shows mean expression, Log2 fold-change (FC ) of *ICN1-GFP*$^{Con}$*/CD4-Cre* and FDR-adjusted *q* values for the indicated probe sets in DP thymocytes. Those highlighted in green had Log2 FC≥0.7 and FDR≤0.05

**Supplementary Table 2: Lower expression of cell cycle, DNA replication and repair genes in *ICN1-GFP*$^{Con}$ DP thymocytes**

| | | | | Mean Expression in DP | | | |
|---|---|---|---|---|---|---|---|
| **Probe_Id** | **Symbol** | **Gene Name** | **Entrez Gene ID** | ***CD4-Cre*** | ***ICN1-GFP*$^{Con}$** | **Log2 FC** | ***FDR*** |
| ILMN_1245789 | *Ccne2* | cyclin E2 | 12448 | 9.39 | 8.14 | -1.24 | 2.25E-03 |
| ILMN_2637704 | *Cdc25c* | cell division cycle 25 homolog C (S. pombe) | 995 | 8.35 | 7.57 | -0.78 | 5.54E-02 |
| ILMN_2919433 | *Cdc45* | cell division cycle 45 homolog (S. cerevisiae) | 12544 | 9.78 | 8.75 | -1.03 | 1.61E-02 |
| ILMN_2597255 | *Cdc6* | cell division cycle 6 homolog (S. cerevisiae) | 23834 | 9.57 | 8.42 | -1.15 | 9.35E-03 |
| ILMN_1238374 | *Cdc7* | cell division cycle 7 (S. cerevisiae) | 12545 | 10.66 | 9.83 | -0.84 | 4.39E-04 |
| ILMN_2832219 | *Cdca3* | cell division cycle associated 3 | 14793 | 10.36 | 9.04 | -1.32 | 1.85E-03 |
| ILMN_2730463 | *Cdt1* | chromatin licensing and DNA replication factor 1 | 67177 | 11.36 | 10.44 | -0.92 | 3.71E-02 |
| ILMN_1236574 | *Cenpa* | centromere protein A | 12615 | 13.84 | 12.96 | -0.88 | 2.80E-02 |
| ILMN_2693785 | *Cenph* | centromere protein H | 26886 | 7.82 | 7.02 | -0.79 | 7.82E-03 |
| ILMN_2821788 | *Cenpi* | centromere protein I | 102920 | 8.42 | 7.71 | -0.71 | 4.06E-02 |
| ILMN_3104928 | *Cenpm* | centromere protein M | 66570 | 8.78 | 7.90 | -0.88 | 2.33E-03 |
| ILMN_1223979 | *Cenpn* | centromere protein N | 72155 | 9.31 | 8.23 | -1.08 | 4.11E-04 |
| ILMN_2970623 | *Cenpp* | centromere protein P | 66336 | 9.81 | 8.73 | -1.08 | 3.09E-02 |
| ILMN_1232175 | *Chtf18* | CTF18, chromosome transmission fidelity factor 18 homolog (S. cerevisiae) | 214901 | 8.82 | 7.93 | -0.89 | 2.20E-02 |
| ILMN_2858359 | *Clspn* | claspin homolog (Xenopus laevis) | 269582 | 10.99 | 9.59 | -1.40 | 2.96E-04 |
| ILMN_2761852 | *D2Ertd750e* | DNA segment, Chr 2, ERATO Doi 750, expressed | 51944 | 8.36 | 7.66 | -0.70 | 1.58E-02 |
| ILMN_2628708 | *Depdc1b* | DEP domain containing 1B | 218581 | 9.70 | 8.76 | -0.93 | 5.36E-03 |
| ILMN_2720451 | *Dnmt1* | DNA methyltransferase (cytosine-5) 1 | 13433 | 10.69 | 9.98 | -0.71 | 2.17E-02 |
| ILMN_1227159 | *Dut* | deoxyuridine triphosphatase | 110074 | 8.49 | 7.77 | -0.73 | 4.00E-02 |
| ILMN_1231138 | *E2f1* | E2F transcription factor 1 | 13555 | 9.53 | 8.60 | -0.93 | 2.97E-03 |
| ILMN_2673776 | *E2f2* | E2F transcription factor 2 | 242705 | 12.68 | 11.49 | -1.18 | 8.21E-04 |
| ILMN_2855315 | *Hist1h1c* | histone cluster 1, H1c | 50708 | 10.68 | 9.91 | -0.77 | 2.38E-06 |
| ILMN_1242399 | *Hist1h2be* | histone cluster 1, H2be | 68024 | 9.23 | 8.32 | -0.91 | 5.00E-04 |
| ILMN_1222313 | *Hist1h4f* | histone cluster 1, H4f | 319157 | 10.38 | 9.65 | -0.73 | 3.27E-02 |
| ILMN_2626619 | *Hist12h4* | histone cluster 1, H4f | 97122 | 8.40 | 7.70 | -0.70 | 5.12E-04 |
| ILMN_2785454 | *Hist2h2ab* | histone cluster 2, H2ab | 621893 | 12.77 | 11.24 | -1.54 | 1.64E-02 |
| ILMN_1231066 | *Hist2h2be* | histone cluster 2, H2be | 319190 | 9.44 | 8.71 | -0.73 | 3.26E-04 |
| ILMN_2866970 | *Kif11* | kinesin family member 11 | 16551 | 10.57 | 9.51 | -1.06 | 4.67E-02 |
| ILMN_2923463 | *Kif23* | kinesin family member 23 | 71819 | 10.25 | 9.52 | -0.73 | 2.28E-04 |
| ILMN_2728300 | *Mdm1* | transformed mouse 3T3 cell double minute 1 | 17245 | 7.98 | 7.26 | -0.72 | 1.15E-04 |
| ILMN_2762026 | *Pcna* | proliferating cell nuclear antigen | 18538 | 8.79 | 8.06 | -0.73 | 4.76E-02 |
| ILMN_2600004 | *Plk4* | polo-like kinase 4 (Drosophila) | 10733 | 8.41 | 7.57 | -0.84 | 4.49E-03 |
| ILMN_2614304 | *Pola2* | polymerase (DNA directed), alpha 2 | 18969 | 9.40 | 8.69 | -0.71 | 1.03E-02 |
| ILMN_2598548 | *Pole* | polymerase (DNA directed), epsilon | 18973 | 8.21 | 7.41 | -0.81 | 9.45E-03 |
| ILMN_1256883 | *Rad51* | RAD51 homolog (S. cerevisiae) | 19361 | 8.35 | 7.51 | -0.84 | 4.78E-02 |
| ILMN_2753226 | *Rad51ap1* | RAD51 associated protein 1 | 10635 | 8.69 | 7.90 | -0.79 | 2.12E-02 |
| ILMN_1239806 | *Rpa3* | replication protein A3 | 68240 | 12.28 | 11.38 | -0.89 | 8.60E-06 |
| ILMN_2613118 | *Rfc4* | replication factor C (activator 1) 4 | 5984 | 10.43 | 9.44 | -0.99 | 2.36E-02 |
| ILMN_2633697 | *Skp2* | S-phase kinase-associated protein 2 (p45) | 27401 | 8.33 | 7.54 | -0.80 | 2.49E-03 |
| ILMN_2615709 | *Spc24* | SPC24, NDC80 kinetochore complex component, homolog (S. cerevisiae) | 67629 | 9.89 | 8.86 | -1.03 | 1.95E-02 |
| ILMN_1255960 | *Spc25* | SPC25, NDC80 kinetochore complex component, homolog (S. cerevisiae) | 66442 | 10.22 | 9.25 | -0.97 | 7.14E-03 |
| ILMN_2893926 | *Tk1* | thymidine kinase 1 | 21877 | 8.20 | 7.45 | -0.75 | 1.72E-02 |
| ILMN_2830661 | *Top2a* | topoisomerase (DNA) II alpha | 21973 | 12.48 | 11.48 | -1.00 | 2.38E-02 |
| ILMN_2416145 | *Tyms* | thymidylate synthase | 22171 | 9.95 | 9.02 | -0.93 | 4.33E-02 |
| ILMN_3150811 | *Ybx2* | Y box protein 2 | 53422 | 7.96 | 7.19 | -0.76 | 1.14E-02 |

*Legend:* Probe sets specific for cell cycle, DNA replication and repair genes that were expressed at significantly lower levels in DP thymocytes from *ICN1-GFP*$^{Con}$ mice. These were selected from 117 expressed at significantly lower levels in mutant DP thymocytes

**Supplementary Table 3: Cell cycle, DNA replication and repair gene sets enriched i**

| *Gene Set Name* | *No.Genes* | *NES* | *FDR* |
|---|---|---|---|
| *REACTOME_CELL_CYCLE* | 421 | -2.68 | 0 |
| *REACTOME_CELL_CYCLE_CHECKPOINTS* | 205 | -2.58 | 0 |
| *REACTOME_DNA_DOUBLE_STRAND_BREAK_REPAIR* | 103 | -2.41 | 0 |
| *REACTOME_DNA_REPAIR* | 218 | -2.40 | 0 |
| *REACTOME_DNA_REPLICATION* | 107 | -2.40 | 0 |
| *REACTOME_MITOTIC_METAPHASE_AND_ANAPHASE* | 182 | -2.38 | 0 |
| *REACTOME_G2_M_CHECKPOINTS* | 106 | -2.18 | 0 |

***Legend*:** GSEA analysis of global gene expression *CD4-Cre* vs *ICN1-GFP*$^{Con}$ DP thymocytes. The table displays gene sets with highly significant negative normalized enrichment scores (NES), indicating significant enrichment for these gene sets in *CD4-Cre* DP thymocytes.

**Supplementary Table 4: Differrential expression of selected genes in *ICN1-GFP$^{Con}$* DP thymocytes**

| Probe_Id | Symbol | Gene Name | Entrez Gene ID | Mean Expression in DP thymocytes (Log2) | | Log2 FC | *FDR* |
|---|---|---|---|---|---|---|---|
| | | | | *CD4-Cre* | *ICN1-GFP$^{Con}$* | | |
| ***A. Activation and Selection*** | | | | | | | |
| ILMN_2821118 | *Ccr7* | *chemokine (C-C motif) receptor 7* | 12775 | 7.05 | 7.19 | 0.14 | |
| ILMN_2659739 | *Il7r* | *interleukin 7 receptor* | 16197 | 6.72 | 6.93 | 0.21 | |
| ILMN_2590211 | *Itm2a* | *integral membrane protein 2A* | 16431 | 6.56 | 6.54 | -0.02 | |
| ILMN_2589871 | *Cd28* | *CD28 antigen* | 12487 | 11.9 | 11.88 | -0.02 | |
| ILMN_2643910 | *Cd5* | *CD5 antigen* | 100045607 | 10.88 | 10.95 | 0.07 | |
| ILMN_1218240 | *Cd69* | *CD69 antigen* | 12515 | 10.52 | 10.95 | 0.43 | |
| ***B. CD4 Lineage*** | | | | | | | |
| ILMN_1249663 | *Cd4* | *CdD4 antigen* | 12504 | 9.26 | 9.45 | 0.20 | |
| ILMN_2441391 | *Zbtb7b* | *zinc finger and BTB domain containing 7B (ThPOK)* | 22724 | 6.85 | 6.86 | 0.01 | |
| ILMN_1257241 | *Cd40lg* | *CD40 ligand* | 21947 | 6.64 | 6.64 | 0 | |
| ***C. CD8 Lineage*** | | | | | | | |
| ILMN_2935080 | *Cd8a* | *CD8 subunit alpha* | 12525 | 6.57 | 6.48 | -0.09 | |
| ILMN_2747141 | *Runx3* | *runt related transcription factor 3* | 12399 | 6.46 | 6.45 | -0.01 | |
| ILMN_2707181 | *Cd160* | *CD160 antigen* | 54215 | 7.06 | 10.27 | 3.21 | 5.50E-11 |
| ILMN_1244891 | *Cst7* | *cystatin F (leukocystatin)* | 13011 | 6.85 | 7.94 | 1.09 | 3.78E-08 |
| ILMN_2699898 | *Itgae* | *integrin alpha E, epithelial-associated* | 16407 | 7.06 | 8.55 | 1.49 | 4.95E-04 |
| ILMN_1228333 | *Prf1* | *perforin 1 (pore formaing protein)* | 18646 | 6.97 | 7.90 | 0.93 | 8.44E-09 |
| ILMN_1217855 | *Nkg7* | *natural killer cell group 7 sequence* | 72310 | 7.18 | 9.95 | 2.77 | 2.78E-08 |
| ILMN_2948552 | *Xcl1* | *chemokine (X-C motif) ligand 1* | 16963 | 6.7 | 8.32 | 1.62 | 8.86E-04 |

*Legend:* The table shows mean expression, Log2 FC (ICN1-GFP$^{Con}$/*CD4-Cre)* and FDR-adjusted *q* values for the indicated probe sets. *Highly significant CD8 lineage genes with FDR<10-3 are highlighed in green.*
